# Experimental evidence that female rhesus macaques (*Macaca mulatta*) perceive variation in male facial masculinity

**DOI:** 10.1101/222810

**Authors:** Kevin A. Rosenfield, Stuart Semple, Alexander V. Georgiev, Dario Maestripieri, James P. Higham, Constance Dubuc

**Affiliations:** Centre for Research in Evolutionary, Social and Interdisciplinary Anthropology, University of Roehampton, Holybourne Avenue, London SW15 4JD, UK; Department of Anthropology, Pennsylvania State University, 409 Carpenter Building, University Park, PA 16802, USA; School of Natural Sciences, Bangor University, Bangor, Gwynedd, LL57 2UW, UK; Institute for Mind and Biology, The University of Chicago, 940 East 57th St, Chicago, IL 60637, USA; Department of Anthropology, New York University, 25 Waverly Place, New York, NY 10003, USA; Department of Zoology, University of Cambridge, Downing Street, Cambridge CB2 3EJ, UK

**Keywords:** Sexual dimorphism, mate choice, facial masculinity, look-time experiment

## Abstract

Among many primate species, face shape is sexually dimorphic, and male facial masculinity has been proposed to influence female mate choice and male-male competition by signalling competitive ability. However, whether conspecifics pay attention to facial masculinity has only been assessed in humans. In a study of free-ranging rhesus macaques, *Macaca mulatta*, we used a two-alternative look-time experiment to test whether females perceive male facial masculinity. We presented 107 females with pairs of images of male faces – one with a more masculine shape and one more feminine – and recorded their looking behaviour. Females looked at the masculine face longer than at the feminine face in more trials than predicted by chance. Although there was no overall difference in average look-time between masculine and feminine faces across all trials, females looked significantly longer at masculine faces in a subset of trials for which the within-pair difference in masculinity was most pronounced. Additionally, the proportion of time subjects looked toward the masculine face increased as the within-pair difference in masculinity increased. This study provides evidence that female macaques perceive variation in male facial shape, a necessary condition for intersexual selection to operate on such a trait. It also highlights the potential impact of perceptual thresholds on look-time experiments.

## Introduction

Sexual selection can shape the evolution of male secondary sex characters through the processes of intra- or intersexual selection, commonly associated with male-male contest competition and female mate choice, respectively [1]. Although intra-sexual and intersexual selection were initially believed to be independent evolutionary processes [1], a growing body of evidence now indicates that traits initially shaped by intrasexual selection - such as badges of dominance status, agonistic displays, large body size and weapons - can sometimes be used secondarily by females as cues or signals of male physical strength and competitive ability, allowing them to select optimal mating partners or avoid coercive males [2–4]. As long as inter-male variation in such traits can be perceived, females might be able to use them in their mating decisions.

In humans, there is good evidence that facial masculinity is associated with male-male competition: facial masculinity has been found to be positively associated with physical strength [5], testosterone levels [6,7; but see 8, in which no link was found, 9, in which testosterone reactivity to competition, but not baseline testosterone levels, were related to facial masculinity, and 10, in which neither reactivity nor baseline levels were related to facial masculinity], aggressiveness [11,12], and unethical behaviour (propensity to deceive in negotiation and cheat to increase financial gain) [13]. There is also indirect evidence that facial masculinity predicts fitness, being negatively associated with the probability of dying from contact aggression [14] and positively associated with number of short-term mating partners [15].perceived facial masculinity and dominance are closely linked [5,16], and recent research has shown that humans find viewing male faces rated as dominant as more rewarding, even when ratings of facial attractiveness are statistically controlled [17,18]. Sexually dimorphic face shape is not merely a result of ontogenetic scaling [19], suggesting that it may have been under selection independently of body size. Importantly, variation in facial masculinity is perceived by the human sensory system: it can be used to assess competitive ability [5,16], and more masculine faces appear to be more attractive to women, at least during the fertile phase of the menstrual cycle [5,20,21]. Together, this suggests that in humans, facial masculinity is under either intra- or intersexual selection, or both.

Previous research has shown that primates pay great attention to conspecifics’ faces [22–24]. Facial shape is sexually dimorphic in many primate species (*e.g.* collared mangabeys, *Cercocebus torquatus*: [25]; rhesus macaques, *Macaca mulatta*: [26]; tufted capuchins, *Sapajus apella*: [27,28]; papionins: [29]), and, as in humans, this is not just a consequence of sexual dimorphism in body size [29]. There is evidence that male facial masculinity plays a role in male-male contest competition in tufted capuchins, *Sapajus apella*: in this species, there is a positive association between male facial masculinity (facial width-to height ratio) and both dominance rank [27] and assertiveness [27,30]. Finally, facial masculinity may be associated with greater bite strength in male primates [19]. While there is evidence that other facial features are perceived and used for individual recognition and social decision-making in primates [31–33], whether inter-individual variation in facial masculinity is perceived by conspecifics is unknown.

In this study, we used an experimental approach to investigate whether free-ranging female rhesus macaques perceive variation in male facial masculinity. In this species, sexual dimorphism in facial features [26] may be associated with bite strength [19]; under the assumption that bite strength is associated with success in contest competition and may reflect overall body strength, facial masculinity thus may serve as a cue of male quality or formidability to females. Therefore, we hypothesized that females would show a visual preference for more masculine male faces. Previous research using looking-time experiments has demonstrated that when conspecific faces are presented alongside other types of stimuli, such as seashells or heterospecific faces, primates show a strong conspecific bias [34–39]. To test this hypothesis, we presented adult females with pairs of photographs of faces of adult males, whose facial masculinity we quantified, in order to test two predictions: (1) females will have a higher overall looking time towards the more masculine male face of the pair, and (2) the proportion of time spent looking at the more masculine face will be positively related to the difference in masculinity between the two faces presented.

## Methods

### Study population

We studied rhesus macaques on Cayo Santiago, a 15.2-hectare island 1 km off the eastern coast of Puerto Rico, managed by the Caribbean Primate Research Centre (CPRC) of the University of Puerto Rico. The population of ca. 1,300 - 1,400 macaques living on the island at the time of the study is descended from a group of 409 individuals brought from India in 1938 [40]. Animals are individually recognizable, with tattoos providing a unique ID and ear notches given when they are yearlings. Dates of birth of all animals are available from long-term records.

### Facial sexual dimorphism measurement

To quantify sexual dimorphism in face shape, we measured facial images of male (N=69) and female (N=27) rhesus macaques, collected during the 2012 and 2013 mating season following a previously described method [41]. Multiple images of males were captured in RAW format from 1–3 m away from subjects using a calibrated Canon EOS Rebel T2i camera with an 18-megapixel CMOS APS sensor and an EF-S 55–250 mm f/4–5.6 IS lens. In order to obtain an image of the male looking straight at the camera, we placed a red plastic apple immediately above the camera lens to attract their attention, and collected several images in a row using the burst function, enabling us to select the most forward-facing image from the series. Immediately after the capture of an image, we took a photograph of a colour standard (X-rite ColourChecker passport) placed in the same location and photographed under the same lighting as the subjects (*i.e*., the “sequential method”: [42–45].

For analysis, we chose only images of fully adult males (median age = 9 years; range = 8-16 years; N=69) and females (median age = 9 years; range = 8-14 years; N=27) looking directly towards the camera. For each image, we digitally measured the sizes of eight facial features in GIMP 2013, as depicted in Fig. 1, and scaled the length of each feature by dividing it by head height (hereafter, relative size). We then compared male and female relative feature sizes using either Mann-Whitney U or independent samples t- tests, depending on normality of the data distribution (See Table 1). The relative sizes of five features (lower face height, jaw width, temporalis height, jaw height, nose length) were larger in male faces, while two features (interpupil distance, face width) did not differ significantly between the sexes, and one feature (eye height) was significantly larger in female faces. We then ran a multiple linear regression model with each facial feature as a predictor variable and sex as the independent variable. We saved the unstandardised predicted variables for use as facial masculinity scores for each male and female image. The derived male (mean ± SE = 1.91 ± 0.026) and female (mean ± SE =1.22 ± 0.039) facial masculinity scores differed significantly (Mann-Whitney U = 10.0, P < 0.001).

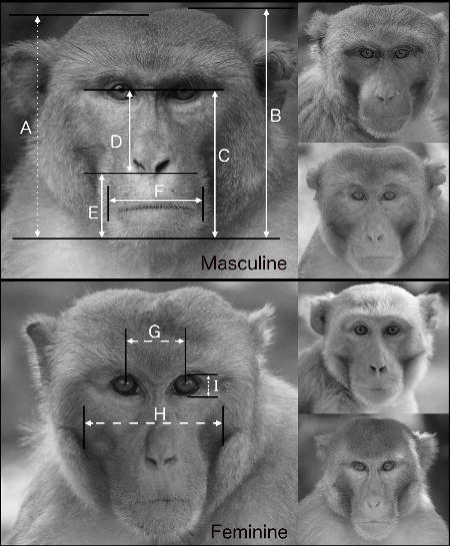

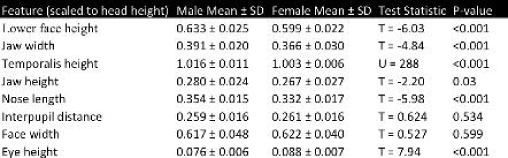

### Stimulus preparation

Using the masculinity scores described above, we selected as stimuli in experimental trials the 10 most masculine and 10 most feminine facial images (hereafter masculine images and feminine images, respectively) that did not contain any distracting elements, such as wounds, discolouration of the facial skin or hair, other monkeys, or food. We only selected images of males displaying neutral expressions, to eliminate the influence of threatening or other facial expressions [46] on subjects’ looking behaviour.

We printed stimuli onto matte photo paper (Staples Photo Supreme) using a colour-calibrated printer (Canon Pixma Pro 100), and measured the printed face colour using a Xrite ColourMunki spectrophotometer (see [31]). Pictures were printed on letter format paper (21.5 cm × 28.9 cm), with printed images of a dimension of 18.5 cm × 18.5 cm, in such a way that face length was 17 cm.

### Experimental design

To test for female preference for male facial masculinity, we used a look-time paradigm that has been used successfully to test interest towards other facial features in this study species [31,46–49]. Each test pair consisted of one masculine and one feminine image, selected randomly from the set of 10 stimuli in each category. KR and one assistant conducted trials on weekdays from 18 March to 29 April 2015, between 09:00 and 13:00 h. The stimuli were placed in frames built into an experimental apparatus, such that they were 85 cm apart at their centres (the apparatus measured 50 x 120 cm; Fig. 2). The relative position of the images in the frame – whether the masculine image was on the right or left – was randomised. Prior to trials, the stimuli were covered by occluders. Potential trial subjects available on Cayo Santiago were all females ≥ 3 years old (N=476 at time of study). We tested 167 of these potential subjects, each being tested only once. We discarded 60 trials that lasted less than 15 s, or during which it was not possible to determine which image the subject was looking toward at any point. This left 107 trials, one from each of 107 subjects (median age = 8 years). Females were not retested if they participated in failed trials, and females that saw stimuli when they were not being tested were also identified and were not tested in future trials. Females who were near adult males, sleeping, or grooming other adults were not tested.

For testing, we placed the experimental apparatus 2-3 m in front of a female and started recording her behaviour on video (Fig. 2). To determine the location of the stimuli in relation to the subject’s eyes (for video coding), we directed her visual attention toward the location of the covered stimuli by tapping on the occluders (in randomised order). We then directed her attention away from either stimulus by tapping on the centre of the apparatus, and removed the occluders to reveal the stimuli. Trials lasted for 30 seconds after removal of the occluders, unless the subject moved away or engaged socially with another monkey. We used MPEG Streamclip for Mac to code only the first 15 seconds (following removal of occluders) of trial videos frame-by-frame, because most subjects stopped looking at either stimulus before the 15^th^ second. During coding, we assessed the total amount of time spent looking at the masculine image and at the feminine image. To eliminate the possibility of coding bias, we coded all trials blind to condition (*i.e.* on which side the masculine image was located).

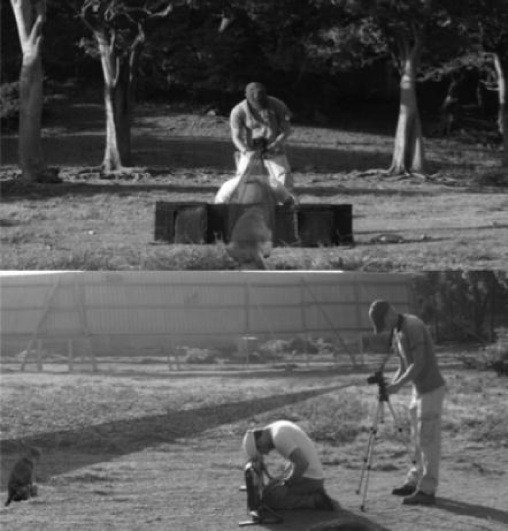

### Potential confounds of facial masculinity

In order to test for possible confounding effects of other male traits, such as age and facial colouration, on stimulus image masculinity, we used Spearman’s rank correlations and Mann-Whitney test. Male age was taken from the long-term records of CRPC. Age and masculinity were not correlated among the males used as experimental stimuli (r_s_ = 0.075, N = 20, P = 0.754), and males used in masculine male images (N = 10, median age = 10) were not older than more feminine males (N = 10; median age = 9; Mann-Whitney U = 39.5, P = 0.421). To quantify facial colour and luminance, we took red (R), green (G) and blue (B) measurements from the stimuli and, based on the processing of colours early in the primate visual pathway, calculated redness as the Red– Green Opponency Channel, (R - G)/(R + G), and darkness as the Luminance (achromatic) Channel (R + G)/2 [46]. Neither facial colour (r_s_ = −0.138, N = 20, P = 0.559) nor facial luminance (r_s_ = 0.339, N = 20, P = 0.143) was correlated with facial masculinity in the stimulus set. Furthermore, masculine (N = 10, median colour = 0.087, median luminance = 167.75) and feminine males (N = 10, median colour = 0.097, median luminance = 145.13) did not differ in facial colour (U = 44, P = 0.684) or luminance (U = 71.5, P = 0.11). We thus concluded that any difference in the looking behaviour of our subjects toward masculine and feminine stimuli would be independent of the effects of male age or facial colour.

We also checked for confounding effects of stimulus males’ familiarity to females. Our operational definition of familiarity was group co-membership. Therefore, using Mann-Whitney tests, we compared the proportion of trial time subjects looked at masculine stimuli when they were groupmates with neither stimulus male (N=76) to trials in which they were groupmates of the masculine stimulus male (N=14), the feminine stimulus male (N=10) and both (N=7). All results were non-significant (masculine vs. neither: U = 543.5, P = 0.903; feminine vs. neither: U = 378, P = 0.983; both vs. neither: U = 345, P = 0.198), indicating that group co-membership did not influence subjects’ looking behaviour.

### Data analysis

To test the prediction that females would look longer at the more masculine face of the pair, we undertook two analyses. First, we compared females’ duration of looking towards the masculine and feminine images using Wilcoxon signed-rank tests. Secondly, we compared the number of trials in which females looked longer at the masculine vs. feminine stimuli to the value expected by chance (0.5) using a binomial test.

We also used two approaches to test the prediction that the proportion of time spent looking at the more masculine face would be positively related to the difference in masculinity between the two faces presented. Firstly, we calculated the relative difference between the masculinity scores of the masculine and feminine image for each trial as follows: ((masculine image score - feminine image score) / feminine image score) x 100, with higher scores indicating larger disparities between the two images. We then ran a linear model with percentage of total look-time spent looking at the masculine image (relative look-time score) as the dependent variable, and relative facial masculinity score as the predictor variable. As relative masculinity may be related to absolute masculinity, we ran an additional linear model, this time entering both relative and absolute facial masculinity score (of the more masculine image) as predictors.

Secondly, to further examine the salience of differences in facial masculinity, we separated the dataset into two groups – one containing the 53 trials with the lowest relative facial masculinity scores, the other containing the 54 trials with the highest scores (results were identical if we used the lowest 54 and the highest 53 trials). We then used Wilcoxon signed-rank tests to compare females’ duration of look-time towards the masculine and feminine images in each group. Statistical tests were two-tailed and performed using IBM SPSS Statistics for Mac (21.0); α was set at 0.05.

## Results

### Prediction 1 - females will look longer at the more masculine male face of the pair

Subjects’ median overall look-time was 4.72 s, or 31% of the 15-s trial period. When all trials were included in the analysis, median looking time for masculine images (2.24 s; IQR = 1.31 – 3.45 s) did not differ from that for feminine images (2.24 s; IQR = 1.07 – 3.43s; Wilcoxon signed-rank test: Z = −0.799, N = 107, P = 0.424; Fig. 3). However, subjects did look longer at the masculine than the feminine image in a significantly higher proportion of trials than expected by chance (looked longer at masculine image: 64 trials; looked longer at feminine image: 41 trials; 2 ties. Binomial test: p = 0.031).

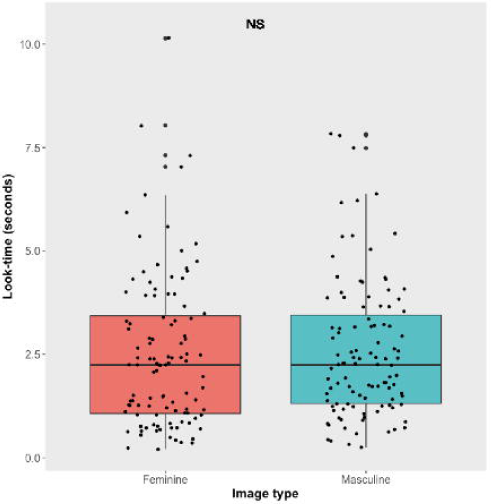

### Prediction 2 - the proportion of time spent looking at the more masculine face will be positively related to the difference in masculinity between the two faces presented

Trials’ relative facial masculinity scores ranged from 12.1% to 61.4%, and variation in these scores explained a small but significant proportion of variation in relative look time scores (β = 0.29, 95% CI =0.03-0.55, p = 0.03, adjusted R_2_ = 0.035; Fig. 4). In other words, the greater the within-trial disparity in masculinity scores, the stronger the bias toward masculine images. The relationship between relative masculinity and relative look time remained significant when absolute masculinity scores were included in the model (β = 0.379, 95% CI =0.06-0.70, p = 0.021), and there was no additional influence of absolute masculinity on relative look times (β = −0.179, 95% CI=0.54-0.192, p = 0.34; total model: F [2,104] = 2.9, adjusted R_2_ = 0.034, p = 0.06).

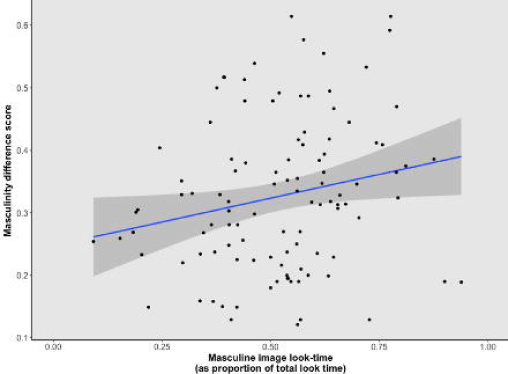

Having separated trials into high (N = 54; mean ± SE = 42.7 ± 0.75%) and low (N = 53; mean ± SE = 22.7 ± 1.14%) relative masculinity score groups, we found that in the low-differences group, subjects’ look-times did not differ between masculine (median = 2.41 s; IQR = 1.48 – 3.66 s) and feminine images (median = 2.41 s; IQR = 1.17 – 3.97 s; Wilcoxon signed-rank test: Z = −1.28, N = 53, P = 0.201; Fig. 5), while in the high-differences group, subjects looked significantly longer at masculine than feminine images (masculine median = 1.86 s; IQR = 1.28 – 2.87 s; feminine median = 1.48 s; IQR = 1.04 – 3.23 s; Wilcoxon signed-rank test: Z = −2.421, N = 54, P = 0.015; Cohen’s d = 0.54; Fig 5).

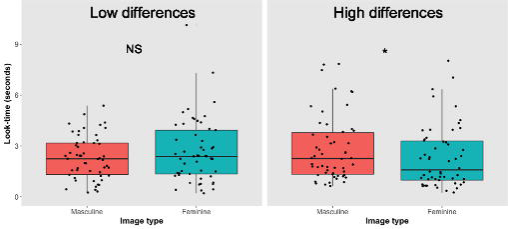

## Discussion

Using a two-alternative experimental look-time paradigm, we tested the hypothesis that free-ranging female rhesus macaques perceive variation in male facial masculinity. In partial support of our prediction that females would look longer at the more masculine male face of the pair, test subjects looked longer at masculine than feminine male faces when the difference in masculinity between the two was high. Moreover, as predicted, the proportion of time spent looking at the more masculine face was positively related to the difference in masculinity between the two faces presented. No relationships were found between male facial masculinity and either male age or facial colour, and female look-time did not appear to be related to familiarity to the test subject, ruling out these as potential confounds of our key results. Overall, this study provides evidence from a non-human species that variation in male facial shape, specifically variation along a feminine-masculine continuum, is salient to female conspecifics.

The finding that females distributed their visual attention unevenly between masculine and feminine faces indicates that the variation in facial masculinity we measured was not only perceived by, but also salient to female rhesus macaques. It is possible that variation in facial masculinity has no reliable connection to underlying physiological, behavioural or genetic factors in male rhesus macaques, in which case there may be no fitness repercussions of female attention to such variation. However, as male facial masculinity is related to hormone levels and behaviour in humans and other primates [*e.g*., 7,43], it seems likely that females’ ability to discriminate variation in this trait is the result of evolutionary processes.

Work to date has indicated that the development of facial masculinity in humans is under the control of testosterone [6,7, but see 10, which failed to replicate this relationship] and is linked to aggressiveness and competitive ability in humans and non-human primates [11,27,30]. As such, it is possible that females gain from paying attention to male facial masculinity because it provides information about the risks of aggression males may present; this explanation has also been proposed to underlie the attentional bias shown for threat grins documented in this species [46,48]. A non-mutually exclusive possibility is that females are attracted to facial masculinity in a sexual context; a preference for males with more masculine faces as mating partners may benefit females if this trait is an honest cue of male genetic quality and health. According to the immuno-competence handicap hypothesis, testosterone-dependent traits can provide information about male quality because androgens are immunosuppressive [51]. Since the development of facial masculinity is under the control of testosterone, high facial masculinity could therefore be a cue to male quality that is available to females. Since visual attentional biases can be underpinned by both attraction and fear (reviewed in [49]), more work is needed to establish whether female perception of variation in male facial features does translate into higher reproductive output for males with more masculine faces, such that female mate choice would play a positive role in maintaining male facial masculinity in this species.

Our finding that subjects’ responses to experimental stimuli depended on relative differences in masculinity highlights the importance of considering aspects of receiver psychology in studies such as this one. We suggest two potential explanations for the positive association between masculinity differences and subjects’ visual bias toward masculine faces. First, the differential responses may have been associated with subjects’ ability to perceive differences in masculinity. A critical feature of signals and cues is that the information they are hypothesised to convey can only alter receiver behaviour if receivers are able to perceive the differences exhibited by emitters [52]; small differences may simply not be discernible. Second, subjects may effectively perceive differences even when small, but such differences may not be sufficient to motivate a differential response; other features, such as skin colouration or texture, may overshadow masculinity differences when they are small.

The effects seen in the present study may represent responses to low-level features (i.e. more elementary features of the scenes presented in our stimuli, such as local colour, luminance or contrast) [53]. In this case, such effects might be seen as the perceptual mechanism by which rhesus macaques are stimulated by masculine facial traits. Such effects would require that low-level features are systematically linked to facial masculinity for them to result in the pattern we observed.

Our study did not attempt to disentangle the potential reasons for the visual biases we observed, but these are important avenues for future investigation. Studies of female rhesus macaque mating behaviour in relation to male characteristics, like those conducted by Manson [54], Dubuc *et al*. [41], and Georgiev *et al*. [55] are needed to determine whether females’ bias in visual attention towards more masculine faces translates into differences in mating and reproductive success. Another important avenue for research is to assess the potential information content of facial shape by investigating the behavioural, physiological, morphological, and genetic correlates of facial masculinity. Finally, as there is evidence that male facial colouration plays an important role in female mate choice in this species [31,41,56], a more comprehensive analysis of the relationship between facial masculinity and facial colouration is needed to better understand how different facial features, and the interaction between them, may shape female preferences.

## Ethics statement

The study was approved by the IACUC of the University of Puerto Rico, Medical Sciences Campus (protocol No. A0100108). All applicable international, national, and/or institutional guidelines for the care and use of animals (in particular non-human primates) were followed. All procedures performed involving animals were in accordance with the ethical standards of the institution(s) at which the study was conducted.

## Data accessibility

All data and code related to this study have been: 1) Submitted as electronic supplementary materials accompanying the manuscript, 2) Submitted to Biorxiv along with manuscript preprint at https://www.biorxiv.org/content/early/2017/11/21/2228103) Uploaded to Dryad data repository; review link:http://datadryad.org/review?doi=doi:10.5061/dryad.k79v6; Temporary Dryad DOI (pending manuscript acceptance): doi:10.5061/dryad.k79v6

## Authors’ contributions

KR conceived, designed, and coordinated the study, measured facial masculinity, performed experiments and statistical analyses, and drafted the manuscript; SS helped design and coordinate the study, carry out statistical analyses, and draft the manuscript; AG and DM helped coordinate the study, and draft the manuscript; JH printed stimulus images, and helped coordinate the study and draft the manuscript; DC collected the stimulus images, and helped design and coordinate the study, carry out statistical analyses, and draft the manuscript. All authors gave final approval for publication.

## Acknowledgements

We would like to thank the staff of the Caribbean Primate Research Centre (CPRC) for assistance in data collection and logistics. Dr. Peter Shaw assisted with the calculation of composite masculinity scores. Sandra Winters helped with image printing at NYU. Field assistance was provided by Severin Mortensen and Maricris Herrera.

